# GOlien tool: fast and scalable GO term annotation of proteins using a Shannon-entropy k-mer model

**DOI:** 10.64898/2026.09.04.749510

**Authors:** Levente Laczkó, Dániel Pék, Béla Lóránt Kovács

## Abstract

**Summary:** Automated protein function annotation remains challenging as sequence databases outpace curated labels and homology-based transfer fails for proteins lacking close relatives. We present GOlien, a scalable, composition-based method that annotates protein sequences using a Shannon-entropy k-mer model. Built from the CAFA3-based training split and evaluated on the held-out validation split, GOlien achieves micro-averaged *F*_*max*_ of 0.7317 (Biological Process), 0.7610 (Cellular Component) and 0.8295 (Molecular Function). Our tool offers a practical, complementary alternative to existing pipelines and extending annotation coverage for proteins.

**Availability and Implementation:** The command line tool to submit FASTA files and retrieve predicted GO term annotations with the corresponding documentation is available at: https://github.com/probalytiq/golien-tool

## Introduction

Predicting the function of (novel) proteins is one of the first tasks in analysing large genomic and proteomic datasets, usually carried out using Gene Ontology (GO) terms (https://geneontology.org/). Although functional annotation of proteins is routinely applied and is an important task influencing the quality of downstream analyses, it is not always a straightforward step. One main reason for inaccurate annotation is the rapid proliferation of protein sequences compared to the slower pace of curated functional annotations; therefore, the generalisation capability of prediction models is crucial.

Approaches include BLAST-based frameworks, which rely on sequence similarity to find the best matching annotation(s) (e.g. BLAST2GO (Conesa *et al*. 2005)), or methods that apply a weighted K-nearest neighbour classifier (e.g. PANNZER2 (Törönen and Holm 2022)). eggNOG-mapper (Cantalapiedra, Hernández-Plaza, Letunic *et al*. 2021) relies on MMseqs2 (Steinegger and Söding 2017), DIAMOND (Buchfink *et al*. 2015) and HMMER3 (Mistry *et al*. 2013) for orthology prediction and protein domain annotation via Pfam. State-of-the-art approaches rely on machine learning methods, such as deep learning (DeepGOPlus (Kulmanov and Hoehndorf 2020)) and specific biological language models (OPUS-GO (Xu *et al*. 2024)), model-attention mechanisms to increase accuracy (PFresGO (Pan *et al*. 2023)), or Siamese neural networks for protein representation (TripletProt (Nourani *et al*. 2022)). The generalisation capability of transformer-based models to unknown proteins could prove beneficial in this field, as implemented in TALE (Cao and Shen 2021) and GOProFormer (Kabir and Shehu 2022), which exploit joint sequence-label embedding strategies. These could be complemented with graph convolutional networks to capture structural variations (Gligorijević *et al*. 2021), or ontology axioms and zero-shot learning to exploit inherent relationships of GO terms (Kulmanov and Hoehndorf 2022).

Homology-based methods, such as BLAST2GO, rely on sequence homology, meaning they primarily identify functions based on closely related proteins. This leads to potential misannotations, especially for proteins with no homologues in the database, resulting in a significant number of unannotated sequences. BLAST2GO also struggles with the resolution of specific functions; it often assigns broader functional classifications instead of more precise ones. Evaluation studies have shown that the metrics used to evaluate PANNZER often favour non-specific and broader classifications over more informative and specific ones, limiting its ability to provide nuanced functional insights (Törönen and Holm 2022).

While neural networks and transformer-based methods show promise in capturing complex relationships between GO terms and improving annotation accuracy, both training and prediction may require significant computational resources and large, high-quality labelled datasets, which may hinder widespread adoption, especially in smaller research settings. In addition, these models may be overfitted to the training data, leading to generalisability issues when new or unrepresented protein sequences are encountered (Oliveira *et al*. 2023). Deep learning models also face challenges related to interpretability, especially in highly complex feature spaces, making it difficult for researchers to understand how specific predictions are made. This lack of explainability, inherent to neural networks used for deep learning, can undermine confidence in these models, particularly in experimental settings where functional predictions guide important decisions (Franchi *et al*. 2022).

In this study, we present GOlien, which annotates protein sequences using a Shannon-entropy k-mer model of protein composition to predict the function of unknown proteins. The platform presented here is readily available and can be applied to support biomedical research by providing high accuracy results with fast running times.

## Methods

The method proposed in this paper annotates protein sequences using a Shannon-entropy k-mer model. Unlike homology-based methods, which transfer annotations from similar sequences, we model sequence composition directly. Each sequence is tokenised into overlapping k-mers, with k-mer size as a tunable hyperparameter, using a rolling-hash representation for efficiency. The Shannon-entropy is then aggregated from the information content of individual k-mers. The range of k-mer sizes (1–5, step size = 1) is chosen to capture both very short and slightly longer local sequence patterns, while keeping the model sparse and computationally manageable.

The training data are taken from the CAFA3-based benchmark dataset of Bianchin de Oliveira *et al*. (2022) (https://doi.org/10.5281/zenodo.7409660). It provides protein sequences with curated Gene Ontology annotations for function prediction evaluation. It is already split into training and validation datasets. For each GO term, we build a k-mer weight map from its annotated training sequences: a mapping from each k-mer to a weight that reflects how characteristic that k-mer is of the term. Our approach can be interpreted as a classification process with two inputs (Figure 1.a). On one side, a reference database of per-GO-term k-mer weight maps is constructed (the model). On the other, the query sequence is tokenised in the same way, using the same k-mer sizes as in training. A query is scored against a GO term by combining the weights its k-mers receive under that term’s weight map, with the score indicating the strength of the match to the term’s k-mer pattern. A query is assigned the GO terms whose scores exceed a calibrated per-term threshold.

**Figure 1.**
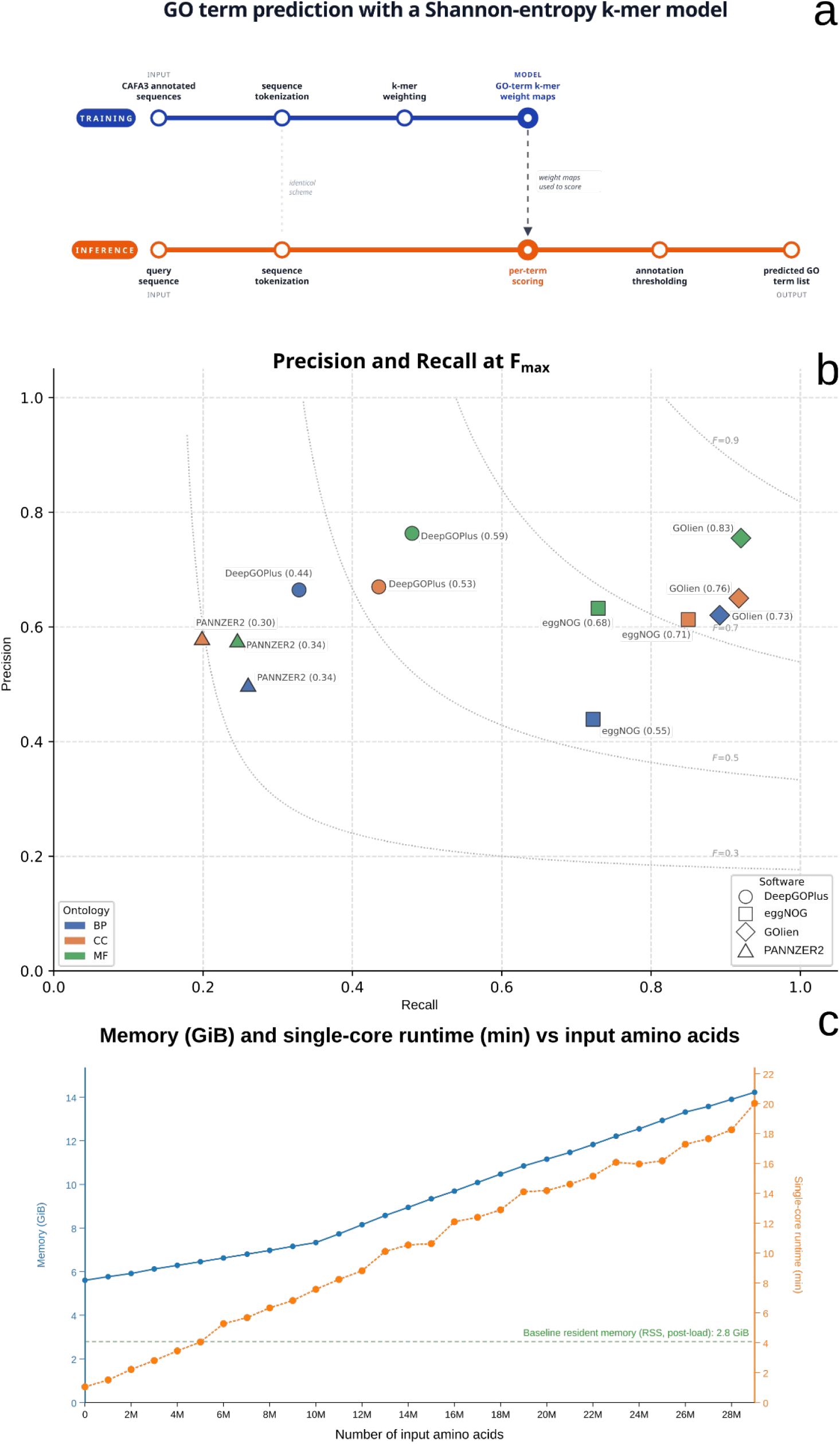
Metro map of the GOlien pipeline. A two-phase pipeline is presented: training derives per-GO-term k-mer weight maps; inference scores query sequences against them. For each GO term, training tokenises its annotated CAFA3 proteins and records a k-mer weight map. Inference tokenises a query, scores it against every weight map, and emits the GO terms whose scores exceed a threshold (a). Relative performance across tools and ontologies, shown as iso- *F*_1_ curves, with each tool’s *F*_*max*_ in parentheses (b). Single-core runtime (minutes, right axis) and peak resident memory (GiB, left axis) versus number of input amino acids, up to ∼2.9 × 10^7^ residue (49,276 sequences); the dashed line marks the post-load model baseline (∼2.8 GiB). Runtime scales linearly; memory grows with input from a fixed ∼5.6 GiB floor reaching ∼14.2 GiB at the largest completed input, near the 16 GB instance limit. (c).

### Validation

The CAFA3-based validation dataset (https://doi.org/10.5281/zenodo.7409660; Bianchin de Oliveira *et al*. 2022) that we used as a gold standard consists of a binary protein × GO term matrix, where each entry indicates whether a given protein is annotated with a given GO term. Evaluations are performed separately for each ontology (Biological Process (BP), Molecular Function (MF), Cellular Component (CC)), using curated term lists to restrict analysis to a well-defined subset of terms. The primary performance metric is the CAFA-style *F*_*max*_ (Radivojac *et al*. 2013), computed as the maximum F1 score over all possible score thresholds.

We selected DeepGOPlus, eggNOG-mapper, and PANNZER2 because they are widely used, publicly available baselines that represent complementary annotation strategies: deep sequence learning, orthology-based annotation transfer, and similarity-based scoring. Together they provide a practical comparison set spanning different approaches. We evaluated all tools on the same CAFA3-based validation dataset to keep the comparison directly comparable.

PANNZER2 produces a continuous ARGOT score for each predicted protein–GO pair, which is used directly as the ranking score. GOlien and DeepGOPlus likewise produce per-prediction scores. For every tool that emits such a score (GOlien, PANNZER2, DeepGOPlus), we sweep the score to trace the full precision–recall curve and report *F*_*max*_ together with the precision and recall at the *F*_*max*_ operating point, so these values are directly comparable. eggNOG-mapper does not provide per-prediction confidence scores in its output format; therefore, each predicted protein–GO pair is assigned a uniform score of 1.0. In this case, *F*_*max*_ reduces to the single operating-point F1 derived from presence/absence predictions. As this corresponds to eggNOG-mapper’s maximum-recall configuration, the comparison is conservative.

For a given score threshold *t*, all predicted (protein, GO term) pairs with a score greater than or equal to *t* are accepted. Precision and recall are then computed over the full set of protein–term pairs jointly across all proteins and all evaluated GO terms:

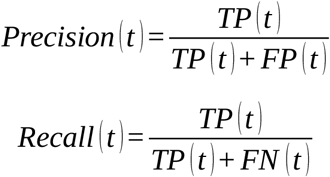

where:

*TP* (*t*) – predicted pairs accepted at threshold that are present in the gold standard,

*FP* (*t*) – predicted pairs accepted at threshold that are absent from the gold standard,

*FN* (*t*) – gold-standard positive pairs not covered by any accepted prediction.

The F1 score at threshold *t* is the harmonic mean of precision and recall:

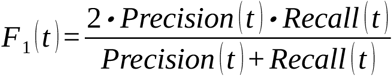

The final reported metric is the maximum F1 over all evaluated thresholds:

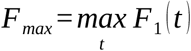

The threshold sweep is implemented by sorting all predictions in descending order of score and updating TP and FP incrementally as each prediction is included. This makes the computation equivalent to tracing the full precision–recall curve without redundant passes over the data.

### Scalability benchmark

To characterise scaling, we ran k-mer counting and inference on increasingly large prefixes of the human reference proteome (GRCh38; RefSeq assembly GCF_000001405.40), taking whole sequences in file order until a target residue budget was reached and stepping the budget in ∼10^6^-amino-acid increments. The largest input that completed within the instance’s memory was ∼2.9 × 10^7^ residues (49,276 sequences); beyond this the process exceeded available RAM. We use the human proteome purely as a large, reproducible source of realistic protein sequences; the scaling behaviour depends only on the number of input residues, not on which proteins are used. Each input size was measured in a separate process to avoid carry-over: the model was loaded once, the resident set after loading recorded as the baseline, and processing then run on a single pinned vCPU with BLAS/OpenMP threads set to one, recording wall-clock runtime and peak resident memory. Benchmarks used a virtualised Linux node (Rackforest (Budapest, Hungary); Intel Xeon Gold host exposed as six single-thread vCPUs under KVM, 16 GB RAM), of which a single vCPU was used.

## Results and discussion

We evaluated GOlien on the CAFA3-based dataset using micro-averaged per-ontology *F*_*max*_ values; the per-ontology *F*_*max*_ and precision/recall at the *F*_*max*_ threshold are as follows: Biological Process (BP) *F*_*max*_ = 0.7317 (precision = 0.6203, recall = 0.8920), Cellular Component (CC) *F*_*max*_ = 0.7610 (precision = 0.6500, recall = 0.9175), and Molecular Function (MF) *F*_*max*_ = 0.8295 (precision = 0.7549, recall = 0.9204).

GOlien outperformed the evaluated baseline methods (DeepGOPlus, eggNOG-mapper, PANNZER2) across all three ontologies (Figure 1.b). The second-best approach – the most competitive on the CC dataset – was eggNOG-mapper (*F*_*max*_ scores: BP = 0.5461, CC = 0.7122, MF = 0.6775), while PANNZER2 and DeepGOPlus yielded *F*_*max*_ scores ranging from 0.30–0.34 and 0.44–0.59, respectively. Relative to these best baselines, GOlien improves *F*_*max*_ by 0.1856 on BP (0.7317 vs 0.5461), 0.0488 on CC (0.7610 vs 0.7122), and 0.1520 on MF (0.8295 vs 0.6775). The observed improvements are most pronounced for BP, suggesting that the k-mer composition signals captured by GOlien are especially informative for broad biological process annotations.

Precision and recall at *F*_*max*_ provide additional context: the evaluation suggests that our approach tends towards high recall at the *F*_*max*_ operating point (recall 0.8920–0.9204 across ontologies) with moderate-to-high precision (0.6203–0.7549) that is comparable to the other baseline methods, indicating that GOlien retrieves a large fraction of true annotations while also emitting some false positives. Some baselines (notably DeepGOPlus) have higher precision at their *F*_*max*_ operating point (e.g. BP precision 0.6645 vs GOlien 0.6203), but GOlien attains markedly higher recall at *F*_*max*_ (BP recall 0.8920 vs best baseline 0.7223; CC recall 0.9175 vs best baseline 0.8498; MF recall 0.9204 vs best baseline 0.7292). Thus, GOlien trades small precision differences for substantially larger recall and overall *F*_*max*_ gains; this trade-off is consistent with a composition-based approach that aggregates many k-mer signals rather than relying solely on strong homology hits.

Overall, the competitors show varying performance across ontologies: eggNOG-mapper is the best baseline on all three ontologies (lowest on BP), DeepGOPlus is comparatively precise but less sensitive, and PANNZER2 shows the lowest overall performance. GOlien appears more balanced across BP, CC, and MF, indicating better cross-ontology generalisation rather than ontology-specific fitting.

GOlien is compact (GO-term k-mer weight maps are stored as serialised sparse representations) and computationally efficient at inference time, making it suitable for large-scale annotation tasks. Because a prediction can be attributed to the k-mers a query shares with a term’s weight map, GOlien is also interpretable: the k-mers most associated with a given GO-term assignment can be identified directly, in contrast to the opaque feature spaces of deep-learning models.

Single-core runtime scaled linearly with input size (Figure 1.c; R^2^ ≈ 0.99, ∼38 s per 10^6^ residues above a fixed per-run cost), reaching ∼20 min for the largest measured input of ∼2.9 × 10^7^ residues (49,276 sequences). Peak memory grew from a fixed floor of ∼5.6 GiB — the ∼2.8 GiB loaded model plus a comparable, input-independent inference working set — to ∼14.2 GiB at that input, at which point the 16 GB instance was exhausted. The fixed cost reflects loading and a single pass over the model that is independent of input, after which both metrics scale with the number of input residues. Because the workload scores each sequence independently, it is trivially parallelisable — expected to scale with the number of available cores — and can be split into chunks to bound peak memory, so inputs larger than a single machine’s RAM, including complete proteomes, can be processed by chunking.

Taken together, GOlien provides a favourable balance of sensitivity and specificity across the evaluated ontologies (particularly on Biological Process and Molecular Function), supporting its use as a scalable approach for large-scale protein function prediction.

## Author contributions

LL: Conceptualization, Validation, Investigation, Visualization, Writing - Original Draft, Writing - Review & Editing; DP: Conceptualization, Methodology, Software, Validation, Investigation, Writing - Review & Editing; BLK: Conceptualization, Methodology, Supervision, Project administration, Writing - Review & Editing

## Conflict of interest

All authors declare that they have no conflicts of interest.

## Funding

None.

## Supplementary Material

**Supplementary Table 1.**
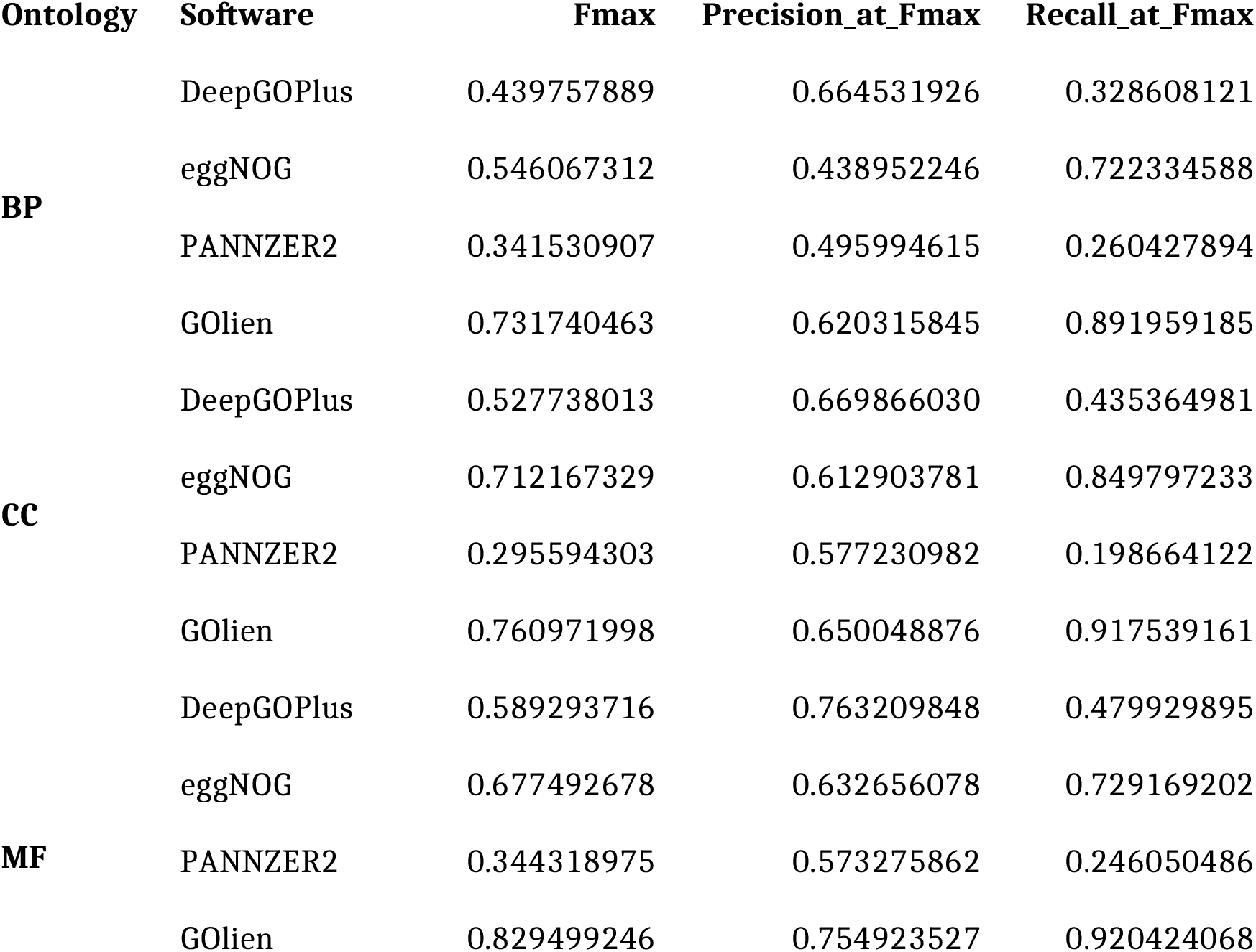
Performance metrics of the baselines and GOlien, micro-averaged per ontology on the CAFA3-based validation split. For eggNOG-mapper, which emits presence/absence calls, *F*_*max*_ reduces to the F1 at its single operating point; precision and recall are those of that point.

| Ontology | Software | Fmax | Precision_at_Fmax | Recall_at_Fmax |
| --- | --- | --- | --- | --- |
| BP | DeepGOPlus | 0.439757889 | 0.664531926 | 0.328608121 |
|  | eggNOG | 0.546067312 | 0.438952246 | 0.722334588 |
|  | PANNZER2 | 0.341530907 | 0.495994615 | 0.260427894 |
|  | GOlien | 0.731740463 | 0.620315845 | 0.891959185 |
| CC | DeepGOPlus | 0.527738013 | 0.669866030 | 0.435364981 |
|  | eggNOG | 0.712167329 | 0.612903781 | 0.849797233 |
|  | PANNZER2 | 0.295594303 | 0.577230982 | 0.198664122 |
|  | GOlien | 0.760971998 | 0.650048876 | 0.917539161 |
| MF | DeepGOPlus | 0.589293716 | 0.763209848 | 0.479929895 |
|  | eggNOG | 0.677492678 | 0.632656078 | 0.729169202 |
|  | PANNZER2 | 0.344318975 | 0.573275862 | 0.246050486 |
|  | GOlien | 0.829499246 | 0.754923527 | 0.920424068 |

## Notes

### Competing Interest Statement

The authors have declared no competing interest.

